# Impacts of retrospective lipid suppression on metabolite quantification in preclinical proton MR spectroscopic imaging

**DOI:** 10.64898/2026.09.14.751356

**Authors:** Tan Toi Phan, Brayan Alves, Bernard Lanz, Cristina Cudalbu

## Abstract

Extracranial lipid contamination remains a challenge in proton magnetic resonance spectroscopic imaging (MRSI), especially in short acquisition delay MRSI, where broad lipid resonances overlap with metabolite and macromolecular signals. Although retrospective lipid suppression techniques are widely used in human MRSI, their effects on metabolite quantification in preclinical MRSI, which is more prone to lipid contamination, have not yet been examined. In this study, we assessed how the retrospective lipid suppression and spectral fitting range influence spectral quality, spatial metabolite mapping, and quantification variability using proton MRSI of rat brains at 14.1 T. Lipid suppression was applied via an orthogonal projection method to both fully sampled and compressed sensing datasets, each comprising data with minimal and pronounced lipid contamination. Spectral fitting was performed with both broad (4.1 - 0.2 ppm) and narrow (4.1 - 1.8 ppm) ranges. When lipid contamination was minimal, suppression caused only slight spectral and spatial changes, with consistent metabolite quantification across conditions. In contrast, datasets with pronounced lipid contamination exhibited notable spectral changes following suppression, consequently affecting spatial metabolite mapping and concentration estimates. Group analysis revealed that metabolites with low concentration estimates were most affected. Similar effects were observed in compressed sensing datasets. Our results provide a better understanding of the impact of retrospective lipid suppression on metabolite quantification in preclinical MRSI, thereby supporting future optimizations for its effective application.

## 1. Introduction

Magnetic resonance spectroscopic imaging (MRSI) has enabled non-invasive, spatially resolved mapping of brain metabolites, providing insights into neuronal integrity, cellular energy metabolism, and membrane turnover^1–3^. Recent advances in pulse sequence design, accelerated acquisition, and spectral-spatial reconstruction have markedly improved spatial resolution and acquisition efficiency, extending the applicability of MRSI in both clinical^4–6^ and preclinical research^7–9^. However, these gains increasingly require data processing pipelines that preserve high spectral fidelity and quantitative accuracy.

A major challenge in quantitative MRSI remains contamination from extracranial and subcutaneous lipids^10,11^. Lipid resonances produce high-amplitude, low-frequency signals that cause spatial ringing, baseline distortions, and spectral overlap with metabolites, most prominently between 1.8 and 0.2 ppm^12–14^. Residual lipid contamination can bias spectral fitting and substantially compromise metabolite quantification, especially in short acquisition delay (AD), short repetition time (TR), fast free-induction decay (FID)-MRSI protocols optimized for maximal signal-to-noise ratio (SNR)^4^.

Prospective lipid suppression strategies, including hardware-based solutions such as crusher coils^15–18^ and sequence-based methods such as saturation bands^19,20^ and chemically selective radiofrequency (RF) pulses^21^ are commonly employed to address this challenge^11^. Sequence-based approaches offer greater flexibility but often yield incomplete suppression in the presence of field inhomogeneities and may inadvertently attenuate cortical metabolite signals, increase specific absorption rate (SAR), and lengthen TR or acquisition time. As a result, prospective suppression alone may be insufficient for robust lipid control.

Retrospective lipid suppression methods have therefore emerged as essential complements to prospective strategies. Techniques such as signal-space projection^12^, low-rank-based lipid basis decomposition^22^, and orthogonal projection^13,23–25^ can substantially reduce lipid contamination.

However, these methods may involve a trade-off between suppression efficacy and the preservation of metabolite signal integrity, which can potentially affect spectral baselines, linewidths, and quantitative accuracy and concentration estimates^6,13,14,26^. This balance is particularly delicate in short AD FID-MRSI, where broad lipid and macromolecular resonances overlap extensively with metabolite peaks, making spectral fitting and metabolite quantification highly sensitive to suppression parameters.

These challenges are further exacerbated in preclinical MRSI. The small nominal voxel size or small acquisition field of view (FOV) required for rodent MRSI reduces SNR and increases susceptibility to both residual lipid contamination and over-suppression. Moreover, rodents are particularly prone to lipid contamination because of the large proportion of surrounding extracranial tissues relative to brain volume. While ultra-high-field systems and cryogenically cooled RF coils partially mitigate SNR limitations, the proximity of brain tissue to extracranial lipid compartments, coupled with increased B_0_ and B_1_ inhomogeneities, makes robust suppression particularly difficult^9,27,28^. Despite a few attempts to use retrospective lipid suppression methods in this setting^29,30^, their quantitative impacts on metabolite concentration estimates in preclinical data have not been systematically evaluated.

Here, we address this gap by evaluating the effect of retrospective lipid suppression (hereafter referred to as lipid suppression) on metabolite quantification in fully sampled and compressed sensing proton FID-MRSI datasets of rat brains acquired at 14.1 T, both with minimal and pronounced lipid contamination. Lipid suppression was performed using a singular value decomposition (SVD)-based orthogonal projection approach^19–21^ implemented in the *MRS4Brain Toolbox*^26,27^. Its impacts were assessed across two spectral fitting ranges: a broad fitting range (4.1 - 0.2 ppm) and a narrow fitting range (4.1 - 1.8 ppm) that excludes lipid-dominated frequencies, thereby allowing evaluation of the influence of a restricted spectral fitting window on metabolite quantification. The analysis focused on N-acetylaspartate (NAA), total choline [tCho = glycerophosphocholine (GPC) + phosphocholine (PCho)], glutamate-glutamine (Glx = Glu + Gln), and myo-inositol (Ins), chosen for their distinct signal amplitudes or concentration estimates, spectral complexity, and varying spectral proximity to lipid resonances. To investigate regional sensitivity to lipid contamination and B_0_ field inhomogeneity associated with reduced shimming quality, concentration estimates for twelve metabolites were analyzed in the hippocampus, striatum, and primary somatosensory cortex. In addition, we analyzed both fully sampled and compressed sensing data^28^. While compressed sensing substantially accelerates MRSI by undersampling *k*-space, it may also alter lipid signal manifestation and spectral characteristics due to acceleration-related artifacts (e.g., lipid-related signal or noise-like aliasing)^8,31^. Therefore, we assessed how the acquisition strategy interacts with lipid suppression in terms of spectral and spatial fidelity, as well as variability in metabolite concentration estimates. Our results improve the understanding of how lipid suppression influences metabolite quantification in preclinical MRSI, thereby supporting future method optimizations.

## 2. Methods

### 2.1. Data acquisition

All experiments were conducted on adult Wistar rats (*n* = 13 rats, approximately 250 ± 50 g, Charles River Laboratories, France). The experimental protocol was approved by the Committee on Animal Experimentation for the Canton of Vaud, Switzerland (VD 3892). The animals were initially anesthetized with 3 - 4% isoflurane in a mixture of oxygen and air (1:1 ratio). They were then placed in a custom-made cradle with head fixation using a pair of ear bars and a bite bar, under anesthesia of 1.5 - 2.5% isoflurane. During the experiment, the animal body temperature was maintained at 37.5°C ± 1.0°C using a warm-water circulation system and measured using a rectal thermosensor. Respiration rate and body temperature were monitored by a small-animal monitor system (SA Instruments, USA).

All scans were performed in a 14.1 T horizontal magnet (Magnex Scientific, UK), interfaced with a Bruker console (ParaVision 360 v1.1 and v3.3), using a custom-made transceiver 2 cm inner-diameter quadrature surface coil (**Fig. 1A**).

**Fig. 1.**
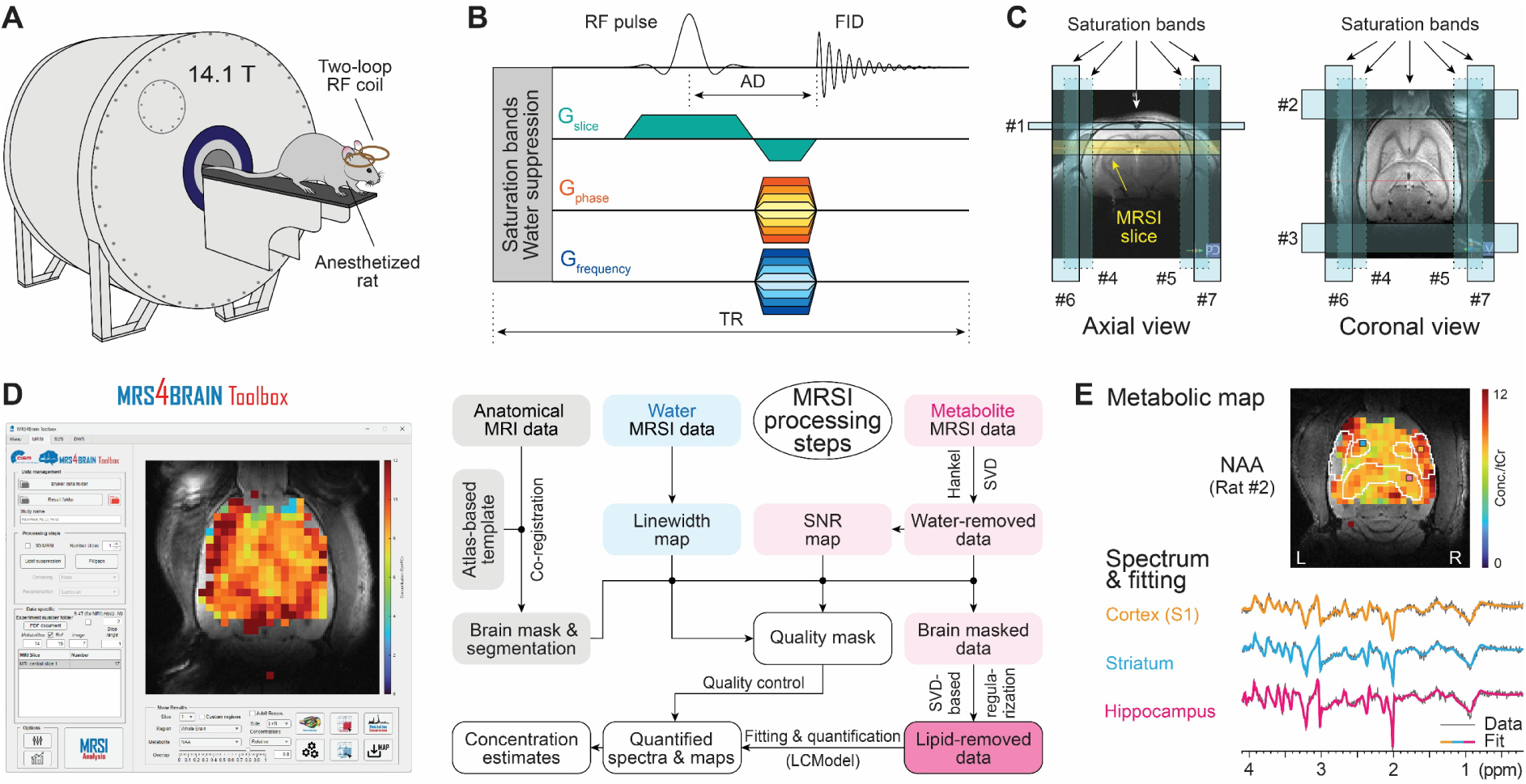
Scheme of proton FID-MRSI data acquisition and analysis in rats at 14.1 T. (**A**) Experiment setup using anesthetized rats in a 14.1 T scanner. (**B**) Schematic drawing of proton FID-MRSI sequence. (**C**) Placement of saturation bands on the rat cranium/skull. (**D**) The *MRS4Brain Toolbox* (left) and its processing pipeline (right) handle all MRSI data analyses, including lipid suppression. (**E**) Representative outputs of the toolbox. Top: NAA metabolite map overlaid on the anatomical image. Bottom: spectra of voxels from the primary somatosensory cortex, striatum, and hippocampus colored in yellow, blue, and pink, respectively. Voxels in the metabolite maps met the quality-control thresholds: SNR ≥ 10, linewidth ≤ 1.25 times the average linewidth, and CRLB < 30%. Metabolite maps are presented relative to tCr fixed at 8 mmol/kgww.

For anatomical reference and brain segmentation, T_2_-weighted Turbo-RARE images were acquired with scan parameters as TR = 4100 ms, echo time (TE) = 27 ms, FOV = 24 × 24 mm^2^, 128 × 128 matrix, 0.188 × 0.188 × 0.2 mm^3^ resolution, 60 coronal slices, and 10 averages.

Proton FID-MRSI data were acquired from a single coronal brain slab, primarily covering the hippocampus, striatum, and primary somatosensory cortex. Each rat underwent both fully sampled and compressed sensing acquisitions. Fully sampled data were acquired using an FID-MRSI sequence (**Fig. 1B**) with the following scan parameters: TR = 811 ms, AD = 1.3 ms, nominal flip angle (FA) = 52°, FOV = 24 × 24 mm^2^, 31 × 31 matrix, nominal voxel size of 0.77 × 0.77 × 2 mm^3^, 7143 Hz spectral bandwidth, and 1024 FID points. For compressed sensing, data were acquired with an acceleration factor of 2 and 20% fully sampled at the *k*-space center. Water suppression was achieved using the variable power RF pulses and optimized relaxation delays (VAPOR) scheme. To reduce extracranial lipid contamination, as rodents are particularly prone to lipid contamination, seven saturation bands were placed in cranial regions (**Fig. 1C**). More detailed experimental and data acquisition information was described in the previous studies^7–9^. Water-unsuppressed data were also acquired for reference using the same parameters, but with the water suppression module disabled. It should be noted that some of the datasets included in this study were used in an earlier work for method demonstration and optimization^8^. However, the analyses presented here are distinct and focus specifically on the effects of lipid suppression, which were not previously investigated.

### 2.2. Data processing

All data were processed using the MATLAB-based *MRS4Brain Toolbox* (MATLAB 2024b), specifically developed for Bruker preclinical proton MRSI data^7,30^. The processing pipeline consists of coregistration of anatomical images with an atlas for brain segmentation, preprocessing steps such as Hankel SVD-based water removal and SVD-based lipid suppression, and fitting and quantification using LCModel^32^ (**Fig. 1D**). The metabolite basis set used for LCModel consisted of 18 metabolites simulated using NMRScopeB/jMRUI, combined with a separately measured *in vivo* macromolecular signal^7^. Each processing step has been explicitly described in previous studies^7,9,30^. Inputs were the reconstructed metabolite and water FID-MRSI data, with Hamming *k*-space filtering, obtained via the vendor reconstruction pipeline for both fully sampled and compressed sensing datasets. Key LCModel fitting parameters were as follows: DKNTMN = 0.25, DEGZER = 0, DEGPPM = 0, SDDEGZ = 999, SDDEGP = 0. We used the total creatine [tCr = creatine (Cr) + phosphocreatine (PCr)] concentration, fixed at 8 mmol/kgww, as an internal reference. Metabolite maps were therefore expressed as relative concentration estimates referring to tCr (hereafter referred to as concentration estimates) and displayed after applying the following quality-control thresholds: SNR ≥ 10, linewidth ≤ 1.25 times the average linewidth of all brain voxels, and Cramer-Rao lower bound (CRLB) ≤ 30% (**Fig. 1E**).

In this study, the lipid suppression algorithm was based on the SVD-based regularization model, described in previous studies^23–25^ and implemented in the *MRS4Brain Toolbox*^7,30^. The method assumes that lipid and metabolite signals are orthogonal in the time or frequency domain, and they do not spatially overlap. First, brain and scalp regions were delineated using the water power mask. Signals from scalp voxels were decomposed via SVD to construct an orthogonal basis representing lipid components. The basis rank was selected by evaluating the energy ratio between brain and scalp regions after applying the corresponding projection operator: E_brain_/E_skull_ ≥ α (default α = 0.8). Consequently, the number of lipid components in the lipid basis was automatically estimated for each rat brain dataset, yielding dataset-specific counts. Datasets with more lipid components in the basis were considered to have more pronounced lipid contamination than those with fewer. Based on this criterion, three datasets from three rats were classified as having pronounced lipid contamination (9 - 63 lipid components), whereas 10 remaining datasets were classified as having minimal lipid contamination (5 - 7 lipid components). The corresponding fully sampled and compressed sensing datasets were consistently assigned to the same group.

To evaluate the influence of residual lipid signals on spectral fitting, we employed two spectral fitting ranges: a broad fitting range (4.1 - 0.2 ppm) and a narrow fitting range (4.1 - 1.8 ppm) that excludes lipid-dominated frequencies. We assessed effects on metabolite-specific fitting variability by analyzing NAA, tCho, Glx, and Ins, selected for their differing signal amplitudes or concentration estimates, complex spectral patterns, and spectral proximity to lipid resonances. To examine sensitivity to lipid source proximity and B_0_ field inhomogeneities associated with reduced shimming quality, we compared averaged metabolite concentration estimates across multiple brain regions, including the hippocampus, striatum, and primary somatosensory cortex. In this analysis, we examined twelve metabolites, including Ins, NAA, taurine (Tau), Glu, total N-acetylaspartate [tNAA = NAA + N-acetylaspartylglutamate (NAAG)], represented for high concentration estimates, and Gln, ascorbate (Asc), aspartate (Asp), gamma-aminobutyric acid (GABA), glutathione (GSH), phosphoethanolamine (PE), and tCho, represented for low concentration estimates. Owing to the ultra-high field strength (14.1 T), all metabolites could be reliably quantified, including metabolites that are difficult to detect because of their low concentration estimates and/or complex spectral patterns (e.g., Asp, GABA, GSH, and PE) or to distinguish due to spectral overlap (e.g., Gln and Glu) at lower field strengths. All analyses were performed on both fully sampled and compressed sensing data.

Coefficient of variation (CV) maps were computed voxel-wise as percentage CV values, calculated as the ratio of the standard deviation to the mean concentration estimate across the pairwise processing conditions being compared. CV values were computed only for spatially corresponding voxels that met the quality-control criteria in both metabolite maps. In other words, we excluded voxels that failed quality control in either metabolite map from CV mapping. For completeness, Glx (Glu + Gln) was included in the CV map analysis as a composite measure of the glutamatergic spectral region. Although Glu and Gln can be reliably quantified separately at 14.1 T, we specifically evaluated Glx because it encompasses a broader spectral region with variable proximity to lipid and macromolecular resonances. This characteristic makes Glx particularly suitable for assessing the effects of lipid suppression on overlapping spectral components and for evaluating the robustness of metabolite quantification in regions susceptible to lipid contamination and macromolecular signals.

Normality of the data was assessed using the Lilliefors test, and homogeneity of variances was evaluated using Levene’s test. When both assumptions were satisfied, a one-way analysis of variance (ANOVA) was conducted. If normality was met but variances were unequal, Welch’s ANOVA was applied. Data that did not fulfill parametric assumptions were analyzed using the Kruskal-Wallis test. Multiple comparisons for all statistical tests were performed using the Bonferroni correction. All data are presented as mean ± standard error of the mean (SEM).

## 3. Results

### 3.1. Impacts on spectral quality and spatial mapping

Proton FID-MRSI protocols with short AD and TR for the rat brain have been extensively optimized in our previous studies^7,9^, resulting in high-quality datasets for this study. All datasets were processed using the same workflow implemented in the *MRS4Brain Toolbox* (**Fig. 1D**), with the same quality control thresholds applied to ensure reliable and consistent quantification results. Consequently, spectra exhibited clearly resolved metabolite peaks across the 4.1 - 0.2 ppm range in different brain regions, and metabolite maps covered the entire slice (**Fig. 1E**). We first evaluated the effects of lipid suppression and spectral fitting ranges (i.e., broad range: 4.1 - 0.2 ppm and narrow range: 4.1 - 1.8 ppm) on spectra acquired under varying levels of lipid contamination in both fully sampled and compressed sensing datasets (**Fig. 2**). This analysis served to characterize case-specific effects before group-level evaluation. In datasets with minimal lipid contamination (**Fig. 2A**), overall spectral quality remained well preserved after lipid suppression across both spectral fitting ranges. Consistent macromolecular fitting (**Fig. S1A**) and small fitting residuals (**Fig. S2A**) were also observed. In contrast, datasets with pronounced lipid contamination exhibited a prominent lipid peak at approximately 1.3 ppm **(Fig. 2B**, top-left). In these cases, lipid suppression substantially reduced lipid contributions, but also aggressively attenuated metabolite and macromolecular signals within the 0.2 - 1.8 ppm range, including partial reduction of the NAA peak near 2 ppm (**Fig. 2B**, top-right). When lipid-suppressed spectra were fitted using the narrow fitting range, spectral mismatches remained visible, particularly around 2 ppm (**Fig. 2B**, bottom-right), similarly to observations obtained with the broad fitting range. The observed mismatches may arise from attenuation of macromolecular and NAA signals following lipid suppression, which may compromise accurate macromolecular fitting, particularly near 0.9 ppm (**Fig. S1B**), thereby contributing to discrepancies in spectral fitting and fitting residuals (**Fig. S2B**). Comparable observations were found in compressed sensing datasets. Lipid suppression resulted in only small spectral alterations in minimally lipid-contaminated spectra (**Figs. 2C**, **S1C**, and **S2C**), specifically near 1.3 - 1.5 ppm in the lipid-suppressed spectrum fitted with the broad range, whereas pronounced lipid contamination led to substantial spectral alterations following lipid suppression (**Fig. 2D**), including compromised macromolecular fitting (**Fig. S1D**) and consequently increased fitting residuals (**Fig. S2D**).

**Fig. 2.**
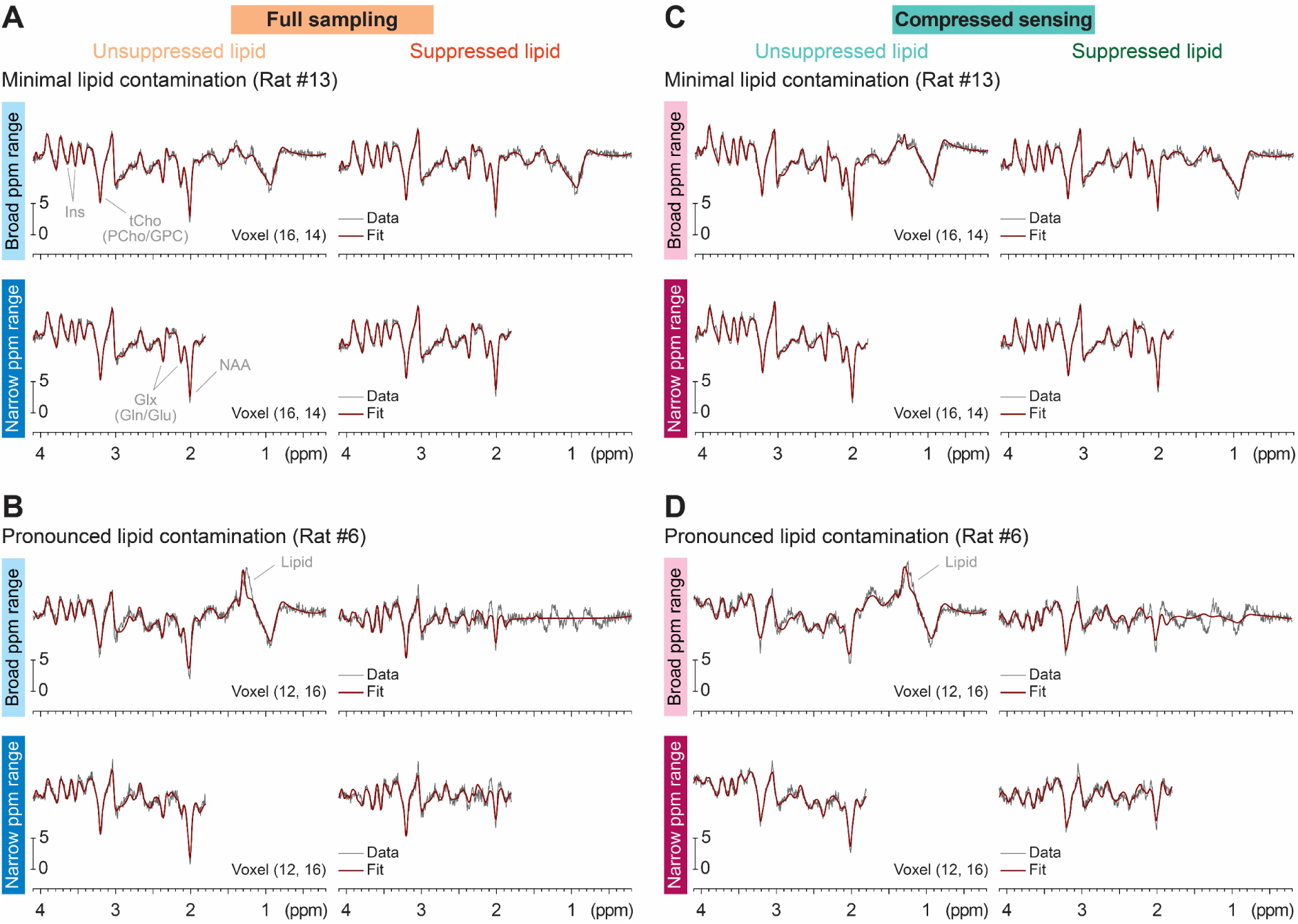
Spectral characteristics under varying suppression and fitting conditions. (**A**) Representative spectra from a fully sampled dataset with minimal lipid contamination (5 lipid components). Top: spectrum fitted over the broad fitting range without (left) and with (right) lipid suppression. Bottom: spectrum fitted over the narrow fitting range without (left) and with (right) lipid suppression. (**B**) Same as (A) but with pronounced lipid contamination (23 lipid components). (**C**) Same as (A) for a compressed sensing dataset (5 lipid components). (**D**) Same as (B) for a compressed sensing dataset (17 lipid components). All signal amplitudes are presented in arbitrary units.

These spectral alterations were further reflected in the spatial distribution and variability of the metabolite maps (**Figs. 3** and **4**). Representative maps of Ins, tCho, Glx, and NAA, selected for their differing signal amplitudes or concentration estimates and spectral patterns in proximity to dominant lipid resonances near 1.3 ppm and macromolecules, were evaluated under varying lipid suppression and fitting conditions. In fully sampled datasets with minimal lipid contamination, metabolite maps remained visually consistent across lipid suppression and spectral fitting ranges (**Fig. 3A**). In datasets with pronounced lipid contamination, there was notable impairment in map appearance following lipid suppression, including altered concentration estimates and reduced spatial coverage for all examined metabolites in both broad and narrow fitting ranges (**Fig. 3C**). These alterations were more evident in tCho and Glx, which exhibit lower concentration estimates and more complex spectral patterns than Ins and NAA. Similar observations were obtained in compressed sensing datasets. Specifically, datasets with minimal lipid contamination showed only small differences across processing conditions (**Fig. 4A**). Conversely, datasets with pronounced lipid contamination demonstrated substantial changes in spatial distribution and map quality following lipid suppression (**Fig. 4C**). However, we observed that compressed sensing datasets exhibited slightly greater changes than fully sampled datasets from the same rat, particularly in brain coverage. This difference may be attributable to a more pronounced manifestation of lipid-related signals in compressed sensing reconstructions, arising from residual spatial aliasing associated with undersampling^31^.

**Fig. 3.**
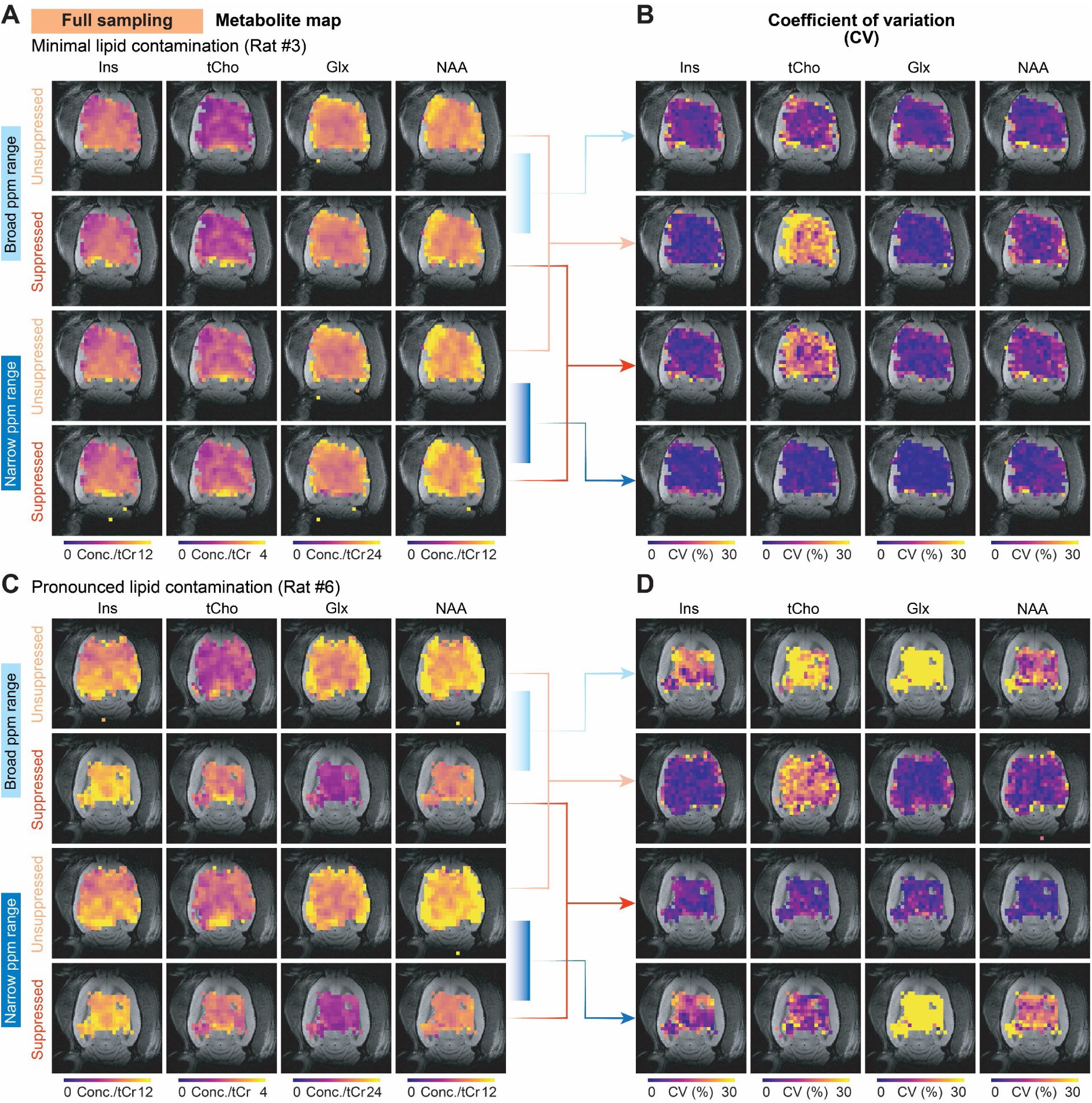
Metabolite and CV maps under varying suppression and fitting conditions for fully sampled data. (**A**) Representative metabolite maps of Ins, tCho, Glx, and NAA from a dataset with minimal lipid contamination (5 lipid components). Rows 1 and 2: maps obtained over the broad fitting range without (row 1) and with (row 2) lipid suppression. Rows 3 and 4: maps obtained over the narrow fitting range without (row 3) and with (row 4) lipid suppression. (**B**) Corresponding CV maps. Rows 1 and 4: comparison between unsuppressed and suppressed conditions in the broad (row 1) and narrow (row 4) fitting ranges. Rows 2 and 3: comparison between broad and narrow fitting ranges under unsuppressed (row 2) and suppressed (row 3) conditions. (**C**) Same as (A) but with pronounced lipid contamination (23 lipid components). (**D**) Same as (B) but with pronounced lipid contamination. Voxels in the metabolite maps met the quality-control thresholds: SNR ≥ 10, linewidth ≤ 1.25 times the average linewidth, and CRLB < 30%. Metabolite maps are presented relative to tCr fixed at 8 mmol/kgww.

**Fig. 4.**
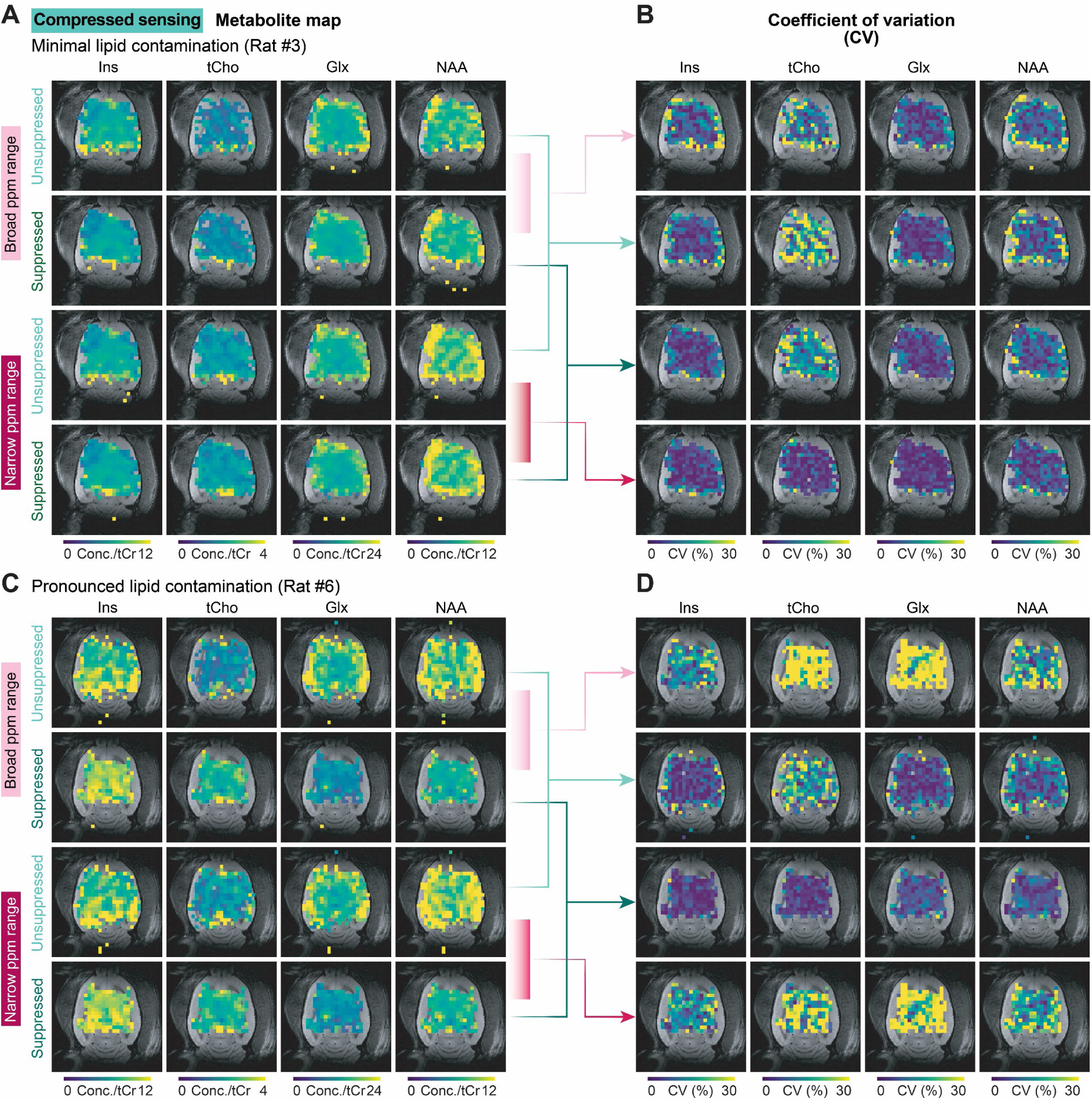
Metabolite and CV maps under varying suppression and fitting conditions for compressed sensing data. (**A**) Representative metabolite maps of Ins, tCho, Glx, and NAA from a dataset with minimal lipid contamination (5 lipid components). Rows 1 and 2: maps obtained over the broad fitting range without (row 1) and with (row 2) lipid suppression. Rows 3 and 4: maps obtained over the narrow fitting range without (row 3) and with (row 4) lipid suppression. (**B**) Corresponding CV maps. Rows 1 and 4: comparison between unsuppressed and suppressed conditions in the broad (row 1) and narrow (row 4) fitting ranges. Rows 2 and 3: comparison between broad and narrow fitting ranges under unsuppressed (row 2) and suppressed (row 3) conditions. (**C**) Same as (A) but with pronounced lipid contamination (17 lipid components). (**D**) Same as (B) but with pronounced lipid contamination. Voxels in the metabolite maps met the quality-control thresholds: SNR ≥ 10, linewidth ≤ 1.25 times the average linewidth, and CRLB < 30%. Metabolite maps are presented relative to tCr fixed at 8 mmol/kgww.

Overall, the effects of lipid suppression depended on the level of lipid contamination. While spectra and metabolite maps were largely preserved in minimally contaminated datasets, pronounced lipid contamination resulted in substantial spectral and spatial alterations following lipid suppression. These effects were further amplified in metabolites having complex spectral patterns and low concentration estimates.

### 3.2. Variability of metabolite fitting

Next, we quantitatively examined variability in metabolite quantification across different lipid suppression and fitting conditions using CV maps. Two representative datasets with minimal and pronounced lipid contamination were used (**Figs. 3** and **4**). For each metabolite, pairwise comparison were performed between processing conditions (**Figs. 3B** and **D**, **4B** and **D**): unsuppressed versus suppressed lipid in the broad fitting range (row 1), unsuppressed versus suppressed lipid in the narrow fitting range (row 4), broad versus narrow fitting ranges under unsuppressed conditions (row 2), and broad versus narrow fitting ranges under suppressed conditions (row 3).

In datasets with minimal lipid contamination, comparisons between metabolite maps processed with and without lipid suppression within the same spectral fitting range showed only minor differences in CV maps (1.75 - 9.46%) across all investigated metabolites, indicating relatively stable metabolite quantification across conditions (**Fig. 3B**, rows 1 and 4). Comparisons between broad and narrow fitting ranges under either unsuppressed or suppressed conditions also exhibited low variability for most metabolites (2.70 - 7.23%), except for tCho (15.04 - 22.69%), which showed greater sensitivity to changes in the spectral fitting range (**Fig. 3B**, rows 2 and 3). By contrast, datasets with pronounced lipid contamination exhibited markedly elevated CV values (11.10 - 49.64%) when comparing unsuppressed with suppressed conditions within the same spectral fitting range (**Fig. 3D**, rows 1 and 4). All metabolites showed increased variability, with tCho and Glx having the highest CV values (11.10 - 29.58% and 45.67 - 49.64%, respectively), indicating greater sensitivity to lipid suppression in the presence of severe lipid contamination. Comparisons between broad and narrow fitting ranges under either unsuppressed or suppressed conditions in this dataset also showed variability similar to those observed in minimally contaminated datasets. CV values remained low for most metabolites in both conditions (2.34 - 6.46%), with tCho exhibiting the greatest variation in the unsuppressed condition (16.92%) (**Fig. 3D**, rows 2 and 3). Similar trends were observed in compressed sensing datasets. (**Figs. 4B** and **D**).

We then extended our observations to the group level (*n* = 13 rats) to provide a more objective evaluation of the effects of lipid suppression and spectral fitting range, using mean CV calculated from all voxels retained in the corresponding CV maps (**Fig. 5**). Overall, the impact of lipid suppression (**Figs. 5A** and **B**, blue and green bars) was significantly greater for the broad fitting range than for the narrow one, particularly for tCho and Glx. In fully sampled data (**Fig. 5A**), mean CV values for tCho were significantly higher in the broad fitting range than in the narrow one (17.72 ± 1.98% versus 7.06 ± 1.07%, *p* < 0.001), with similar findings for Glx (11.32 ± 3.49% versus 8.22 ± 3.42%, *p* < 0.01). Comparable findings were observed in compressed sensing data, where tCho showed CV values of 18.02 ± 2.19% versus 10.56 ± 1.69% (*p* < 0.01), and Glx showed values of 9.31 ± 2.47% versus 6.74 ± 2.28% (*p* < 0.05) for broad and narrow fitting ranges, respectively (**Fig. 5B**). No statistically significant effects of lipid suppression were observed for Ins and NAA.

**Fig. 5.**
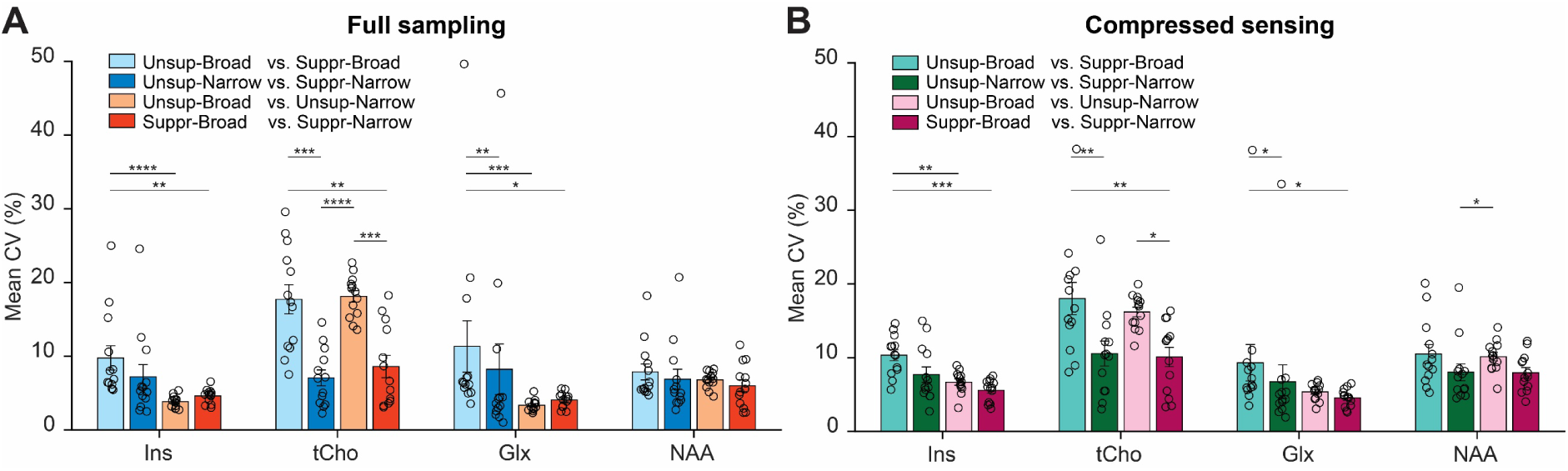
Mean CV under varying suppression and fitting conditions. (**A**) Bar plots of mean CV comparing varying suppression and fitting conditions in Fig. 3 for groups of fully sampled datasets (*n* = 13 rats). (**B**) Bar plots of mean CV comparing varying suppression and fitting conditions in Fig. 4 for groups of compressed sensing datasets (*n* = 13 rats). Mean CV values were calculated from all voxels retained in the corresponding CV maps. Comparisons include unsuppressed versus suppressed lipid in the broad fitting range (light blue or light green), unsuppressed versus suppressed lipid in the narrow fitting range (blue or green), broad versus narrow fitting ranges under unsuppressed conditions (light orange or light pink), and broad versus narrow fitting ranges under suppressed conditions (orange or pink). All data are mean ± SEM. *: *p* < 0.05, **: *p* < 0.01, ***: *p* < 0.001, and ****: *p* < 0.0001 for one-way ANOVA, Welch’s ANOVA, and Kruskal-Wallis tests with Bonferroni multiple comparisons.

When evaluating the effect of spectral fitting range (**Fig. 5**, orange and pink bars), unsuppressed conditions produced greater changes than suppressed conditions, particularly for tCho. In fully sampled data, tCho showed significantly higher CV values without lipid suppression than with lipid suppression (18.12 ± 0.79% versus 8.59 ± 1.51%, *p* < 0.001) (**Fig. 5A**). A similar trend was observed for tCho in compressed sensing data (16.19 ± 0.65% versus 10.09 ± 1.32%, *p* < 0.05) (**Fig. 5B**). No significant effects of spectral fitting range were observed for other examined metabolites.

Overall, these results indicate that metabolite quantification remained stable in minimally contaminated datasets, whereas pronounced lipid contamination increased variability following lipid suppression, particularly for tCho and Glx. These effects were stronger with a broad fitting range and consistent across both fully sampled and compressed sensing data. The observed group-level effects may also have been influenced by datasets with pronounced lipid contamination.

### 3.3. Effects on concentration estimates in different brain regions

Finally, we assessed group-level effects (*n* = 13 rats) on concentration estimates in the hippocampus, striatum, and primary somatosensory cortex, which were influenced by various sensitivity factors such as lipid-source proximity, shimming quality, and field inhomogeneity. Twelve metabolites were classified into two groups: high (Ins, NAA, Tau, Glu, and tNAA) and low (Gln, Asc, Asp, GABA, GSH, PE, and tCho) concentration estimates (**Fig. 6**). In the fully sampled data, we observed that lipid suppression with a broad fitting range (**Figs. 6A - C**, light orange versus orange bars) resulted in statistically significant changes in averaged concentration estimates for tNAA, Gln, and tCho across all brain regions and for GABA in the hippocampus and striatum, compared with the unsuppressed condition. Specifically, the changes for tNAA, Gln, tCho, and GABA amounted to 13.62%, 22.64%, 23.81%, and 36.97%, respectively, in the hippocampus and 13.78%, 22.23%, 22.45%, and 40.25%, respectively, in the striatum, whereas in the primary somatosensory cortex, the corresponding changes were 14.29%, 31.19%, and 20.66% for tNAA, Gln, and tCho, respectively. Lipid suppression with a narrow fitting range affected only GABA (31.39%) in the hippocampus. In compressed sensing data, similar effects were observed in a broad fitting range (**Figs. 6D - F**, light pink versus pink bars), with particularly significant changes for tNAA (11.20%), Gln (16.04%), and tCho (19.41%) in the hippocampus; Gln (15.90%) and tCho (19.58%) in the striatum; and only Gln (20.53%) in the primary somatosensory cortex. There were no effects of lipid suppression within the narrow fitting range in compressed sensing data.

**Fig. 6.**
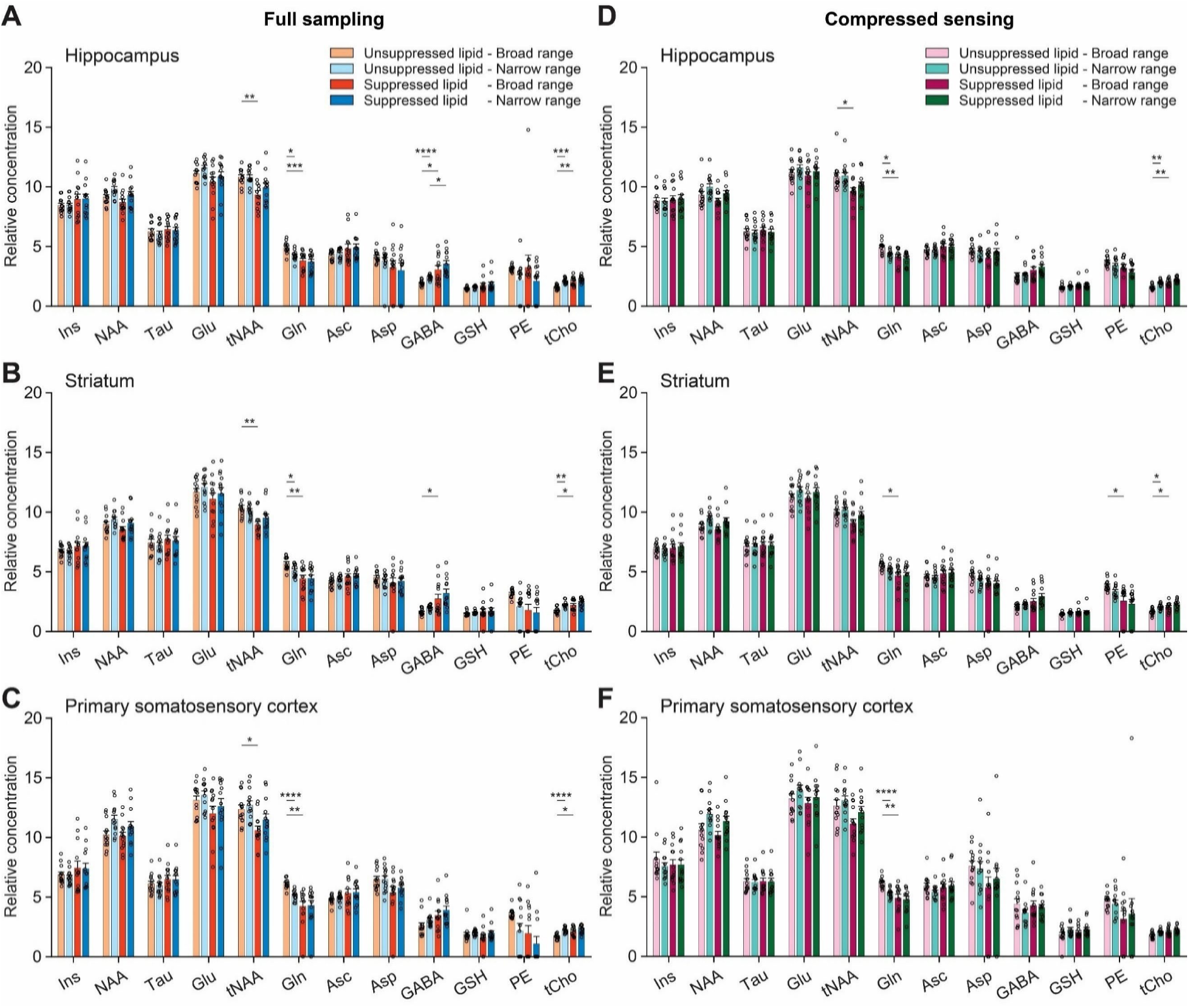
Regional metabolite concentration estimates under varying suppression and fitting conditions. (**A**) Concentration estimates from fully sampled datasets comparing varying suppression and fitting conditions for the hippocampus (*n* = 13 rats). (**B**) Same as (A) but for the striatum (*n* = 13 rats). (**C**) Same as (A) but for the primary somatosensory cortex (*n* = 13 rats). (**D** - **F**) Same as (A - C) for compressed sensing datasets (*n* = 13 rats). All data are mean ± SEM. *: *p* < 0.05, **: *p* < 0.01, ***: *p* < 0.001, and ****: *p* < 0.0001 for one-way ANOVA, Welch’s ANOVA, and Kruskal-Wallis tests with Bonferroni multiple comparisons.

Interestingly, in fully sampled data without lipid suppression, changing the spectral fitting range also led to substantial variations in concentration estimates for Gln and tCho across all brain regions and for GABA in the hippocampus (**Figs. 6A - C**, light orange versus light blue bars). On average, Gln and tCho respectively varied by 24.22% and 30.15% in the hippocampus, 21.74% and 28.56% in the striatum, and 30.10% and 26.85% in the primary somatosensory cortex, whereas GABA changed by 45.42% in the hippocampus when comparing broad versus narrow fitting range. Similar to fully sampled data, where most effects were observed in metabolites with low concentration estimates, compressed sensing data also exhibited a similar effect, with significant changes for Gln (18.79%) and tCho (27.26%) in the hippocampus, only tCho (27.59%) for the striatum, and only Gln (22.66%) for the primary somatosensory cortex (**Figs. 6D - F**, light pink versus light green bars). With lipid suppression, changing the spectral fitting range does not statistically affect concentration estimates for both fully sampled and compressed sensing data (**Figs. 6A - C**, orange versus blue bars; and **Figs. 6D - F**, pink versus green bars).

Additionally, we further examined whether voxels in anatomical regions that met the same quality control thresholds were preserved under the processing conditions. The number of voxels for each processing condition and each brain region was counted. We observed that the number of voxels across processing conditions remained statistically similar for metabolites with high concentration estimates in all examined brain regions, but varied dramatically for metabolites with low concentration estimates, such as Asp, GABA, GSH, and PE (**Figs. 7A - C**). There are similar variations in voxel counts between fully sampled and compressed sensing data (**Figs. 7D - F**). These results further demonstrated that lipid suppression induced spectral alterations when the data were contaminated with lipids, thereby affecting metabolite quantification and concentration estimates, leading to distortions in spatial mapping, especially for metabolites with low concentration estimates.

**Fig. 7.**
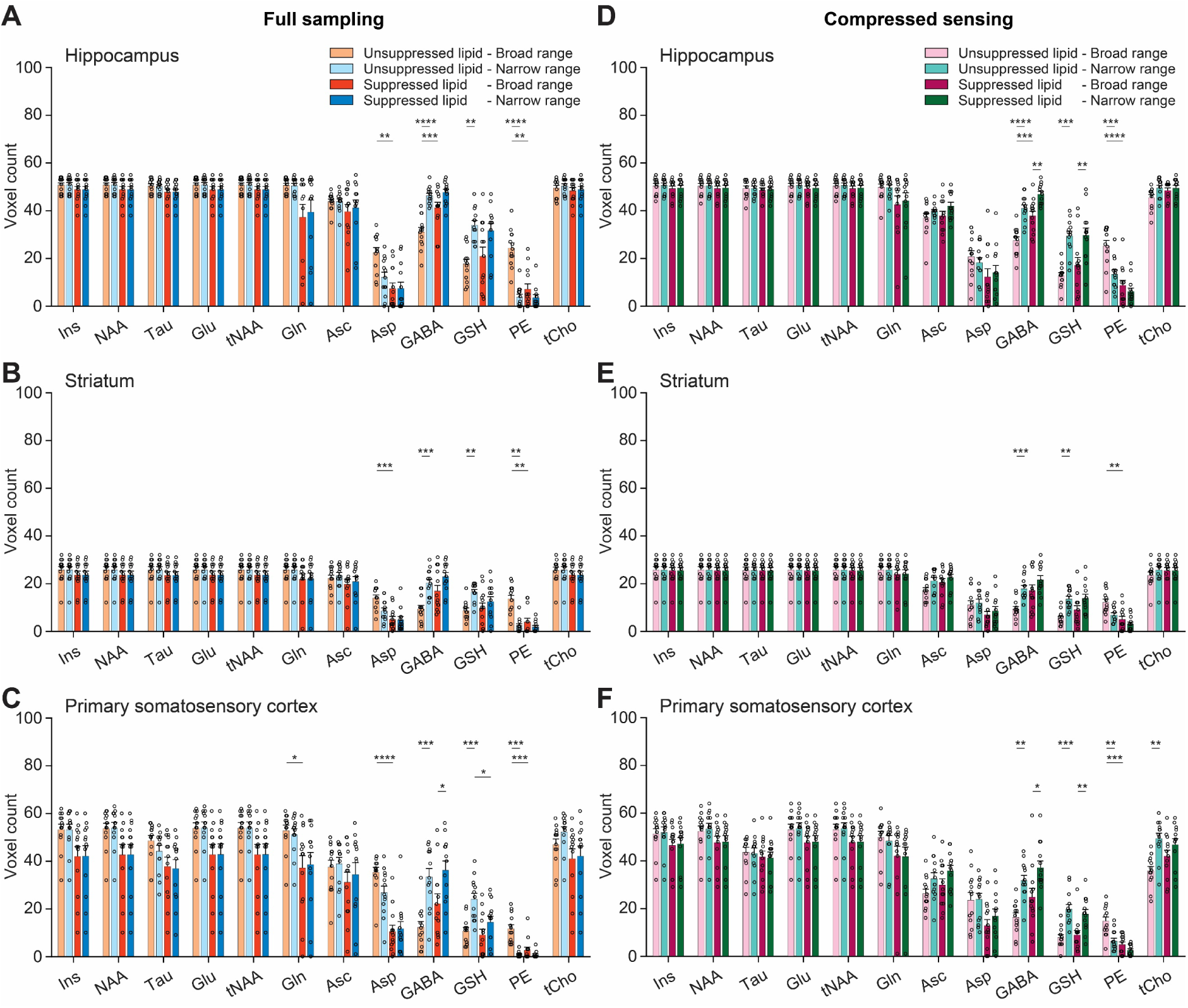
Regional voxel counts in metabolite maps under varying suppression and fitting conditions. (**A**) Regional voxel counts in metabolite maps from fully sampled datasets comparing varying suppression and fitting conditions for the hippocampus (*n* = 13 rats). (**B**) Same as (A) but for the striatum (*n* = 13 rats). (**C**) Same as (A) but for the primary somatosensory cortex (*n* = 13 rats). (**D - F**) Same as (A - C) but for compressed sensing datasets (*n* = 13 rats). All data are mean ± SEM. *: *p* < 0.05, **: *p* < 0.01, ***: *p* < 0.001, and ****: *p* < 0.0001 for one-way ANOVA, Welch’s ANOVA, and Kruskal-Wallis tests with Bonferroni multiple comparisons.

Together, lipid suppression and spectral fitting range primarily affected metabolites with low concentration estimates, notably Gln, GABA, and tCho, with effects varying across brain regions. These processing conditions also influenced voxel preservation and spatial coverage, thereby affecting the robustness of metabolite mapping.

## 4. Discussion

In this study, we systematically examined how lipid suppression and spectral fitting range affect metabolite quantification in preclinical proton FID-MRSI at ultra-high field. Overall, datasets with minimal lipid contamination remained largely stable across processing conditions. In contrast, datasets with pronounced lipid contamination exhibited substantial spectral alterations, increased variability in fitting and quantification, impaired spatial mapping, and reduced voxel retention following lipid suppression^26^. These effects were particularly evident for metabolites with lower concentration estimates or lower signal amplitudes due to J-coupling, such as Gln, GABA, and tCho. They were exacerbated when using broad fitting range that included lipid-dominated frequencies.

The choice of spectral fitting range emerged as a factor influencing these outcomes. Restricting the fitting range, for example, 4.1 - 1.8 ppm, improved fitting and quantification robustness by excluding lipid-dominated frequencies. However, this strategy introduces important trade-offs. Specifically, it prevents fitting and quantification of metabolites resonating within the excluded 0.2 - 1.8 ppm range, such as lactate and alanine. It may also lead to underestimation of macromolecular contributions by LCModel due to the omission of the prominent macromolecular signal near 0.9 ppm.

In addition to extracranial lipid signals, macromolecular resonances represent a further source of complexity^33^. Because these resonances substantially overlap with lipid signals in short AD proton FID-MRSI, lipid suppression may unintentionally attenuate macromolecular components alongside lipids. This likely contributes to less reliable macromolecular estimates, alterations in the spectral baseline, and biased metabolite concentration estimates observed after lipid suppression.

Regional differences further support the role of lipid spatial proximity in driving these effects. Although voxel counts for metabolites with higher concentration estimates in the primary somatosensory cortex did not show statistically significant changes when applying lipid suppression (*p* > 0.05), they exhibited greater variability compared to those in the hippocampus or striatum. This likely reflects the anatomical proximity of cortical regions to the skull and scalp, where extracranial lipid signals are strongest, increasing susceptibility to lipid leakage and suppression-induced distortions. In addition, B_1_ and B_0_ inhomogeneities, as well as poorer shimming in peripheral brain regions that can reduce SNR, may further contribute to voxel loss in the primary somatosensory cortex.

Importantly, similar observations in both fully sampled and compressed sensing datasets indicate that these effects primarily arise from lipid suppression and fitting conditions rather than from the acquisition strategy itself. Although slight differences in brain coverage were observed (**Figs. 3** and **4**), the overall consistency across acquisition approaches strongly supports these findings.

Together, these observations underscore the critical balance between effective lipid removal and the preservation of metabolite signal integrity in preclinical FID-MRSI. While narrowing the spectral fitting range can mitigate lipid-related bias, it inherently limits metabolic coverage and may bias macromolecule estimation. Therefore, lipid suppression should be applied with caution, particularly in datasets with substantial lipid contamination, as overly aggressive processing may compromise the reliability and interpretability of metabolite quantification.

There are several limitations in this work. First, we used single-slice proton FID-MRSI data from rat brains, as developed in our previous studies^7,8^. Multi-slice or three-dimensional (3D) MRSI acquisitions may provide more comprehensive insights into the effects of lipid suppression, as dorsal and ventral regions are likely to experience different levels of lipid contamination^8^. These regions may also vary in coil sensitivity profiles, B_0_ and B_1_ inhomogeneities, shimming quality, and the effectiveness of saturation bands, all of which could influence the observed effects. Second, compressed sensing data used in this study were acquired with a relatively low acceleration factor of 2 and 20% fully sampled at the *k*-space center, which may minimize the effects of lipid suppression on acquisition strategies^8^. Higher acceleration factors have been reported to introduce stronger point-spread function effects and increased lipid-related artifacts^31^, which could further amplify the effect observed here. Lastly, although high-quality proton FID-MRSI datasets acquired with saturation bands were employed to mitigate lipid contamination in this study, subject-specific lipid content can still result in varying degrees of residual lipid contamination across datasets. This issue is particularly pronounced in rodent studies, where the relatively large proportion of extracranial tissues (e.g., skin, muscle, and subcutaneous fat) compared to the small rodent brain volume increases susceptibility to lipid contamination relative to human studies^28^. These considerations underscore the importance of careful optimization and adaptation of saturation band placement and parameters to improve data quality and minimize residual lipid effects^9^.

In future work, we will evaluate the proposed method under more challenging acquisition conditions, including protocols without saturation bands and acquisitions using volume-coil excitation, both of which are expected to increase lipid contamination. The method will also be investigated in 3D MRSI acquisitions, where lipid contamination is typically more pronounced.

In addition, the algorithm’s automatic selection of the number of lipid components will be further refined to improve robustness across different acquisition settings. Finally, alternative lipid suppression approaches, including the water and lipid neural network (WALINET)^14^, will be explored and systematically compared with the proposed framework in this study to assess their relative strengths and limitations.

## 5. Conclusion

In summary, lipid suppression can potentially affect metabolite quantification in preclinical FID-MRSI, especially for metabolites with low concentration estimates. Minimally lipid-contaminated datasets remain relatively stable, whereas pronounced lipid contamination increases fitting and quantification variability. Selecting appropriate lipid suppression and spectral fitting strategies is therefore crucial for preserving signal quality and achieving accurate, reliable metabolite quantification.

## Acknowledgments

We acknowledge the CIBM Center for Biomedical Imaging of the École polytechnique fédérale de Lausanne (EPFL), the Université de Lausanne (UNIL), the Centre Hospitalier Universitaire Vaudois (CHUV), the Université de Genève (UNIGE), and the Hôpitaux Universitaires de Genève (HUG), for providing expertise and resources to conduct this study. We thank Dr. Antoine Klauser for support in implementing the retrospective lipid suppression method in the *MRS4Brain Toolbox*, Drs. Dunja Simicic and Thanh Phong Lê for valuable discussions and support in data acquisition, and the veterinary team for handling the animals.

## Data Availability Statement

The *MRS4Brain Toolbox* and demonstration datasets are publicly available on the GitHub repository^34^ or Zenodo (DOI: 10.5281/zenodo.18270164) or our MRS4Brain group’s website (https://www.epfl.ch/labs/mrs4brain/ressources/mrs4brain-toolbox/).

## Abbreviations

MRSI: Magnetic resonance spectroscopic imaging
AD: Acquisition delay
TR: Repetition time
FID: Free-induction decay
SNR: Signal-to-noise ratio
CRLB: Cramer-Rao lower bound
RF: Radiofrequency
SAR: Specific absorption rate
FOV: Field of view
TE: Echo time
FA: Flip angle
VAPOR: Variable power RF pulses and optimized relaxation delays
SVD: Singular value decomposition
CV: Coefficient of variation
WALINET: Water and lipid neural network
Cr: Creatine
PCr: Phosphocreatine
tCr: Cr + PCr
Glu: Glutamate
Gln: Glutamine
Glx: Glu + Gln
NAA: N-acetylaspartate
NAAG: N-acetylaspartylglutamate
tNAA: NAA + NAAG
GPC: Glycerophosphocholine
PCho: Phosphocholine
tCho: GPC + PCho
Ins: Myo-inositol
Tau: Taurine
Asc: Ascorbate
Asp: Aspartate
GABA: Gamma-aminobutyric acid
GSH: Glutathione
PE: Phosphoethanolamine

