## Supplementary Information for "Impacts of retrospective lipid suppression on metabolite quantification in preclinical proton MR spectroscopic imaging"

**Supplementary Figures:**

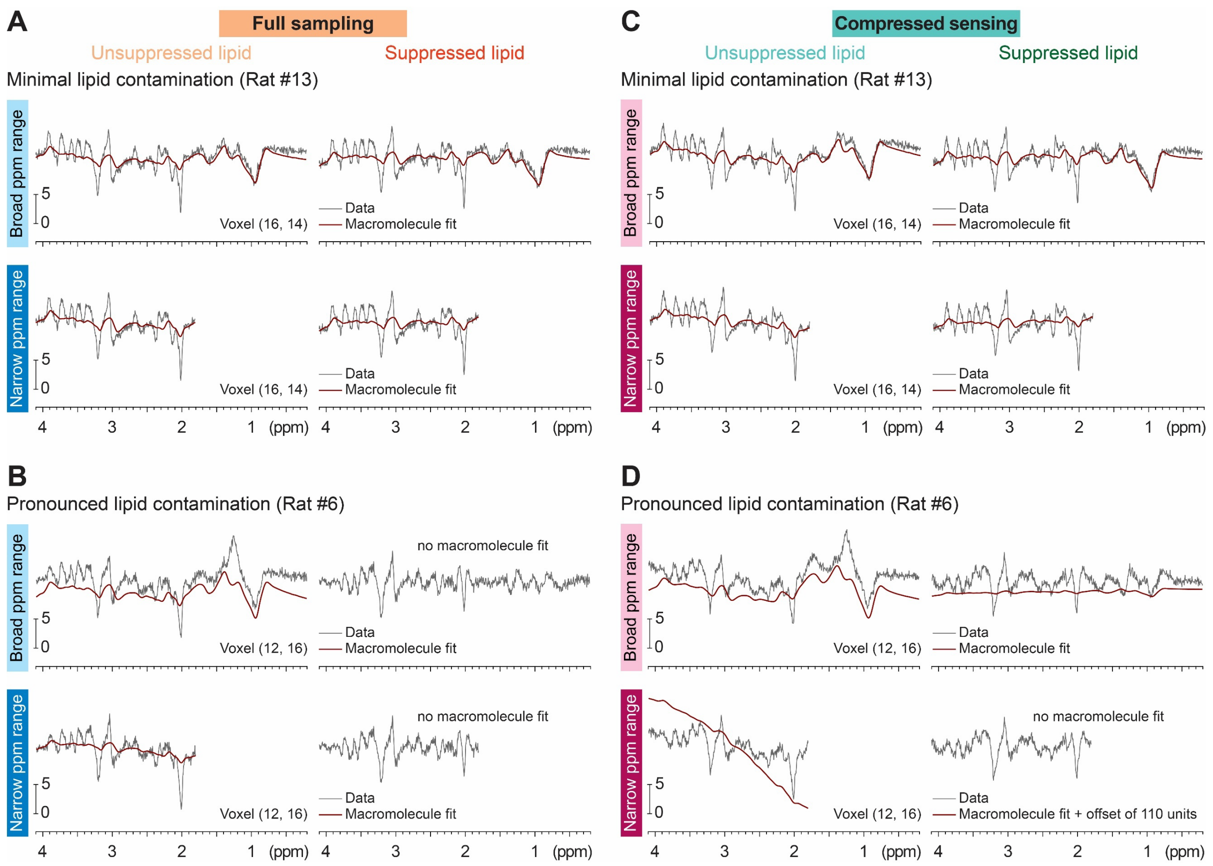


**Fig. S1. Macromolecular fits under varying suppression and fitting conditions in Fig. 2**

(**A**) Macromolecular fits of representative spectra from a fully sampled dataset with minimal lipid contamination (5 lipid components). Top: Macromolecular fits over the broad fitting range without (left) and with (right) lipid suppression. Bottom: Macromolecular fits over the narrow fitting range without (left) and with (right) lipid suppression.

(**B**) Same as (A) but with pronounced lipid contamination (23 lipid components).

(**C**) Same as (A) for a compressed sensing dataset (5 lipid components).

(**D**) Same as (B) for a compressed sensing dataset (17 lipid components).

All signal amplitudes are presented in arbitrary units.

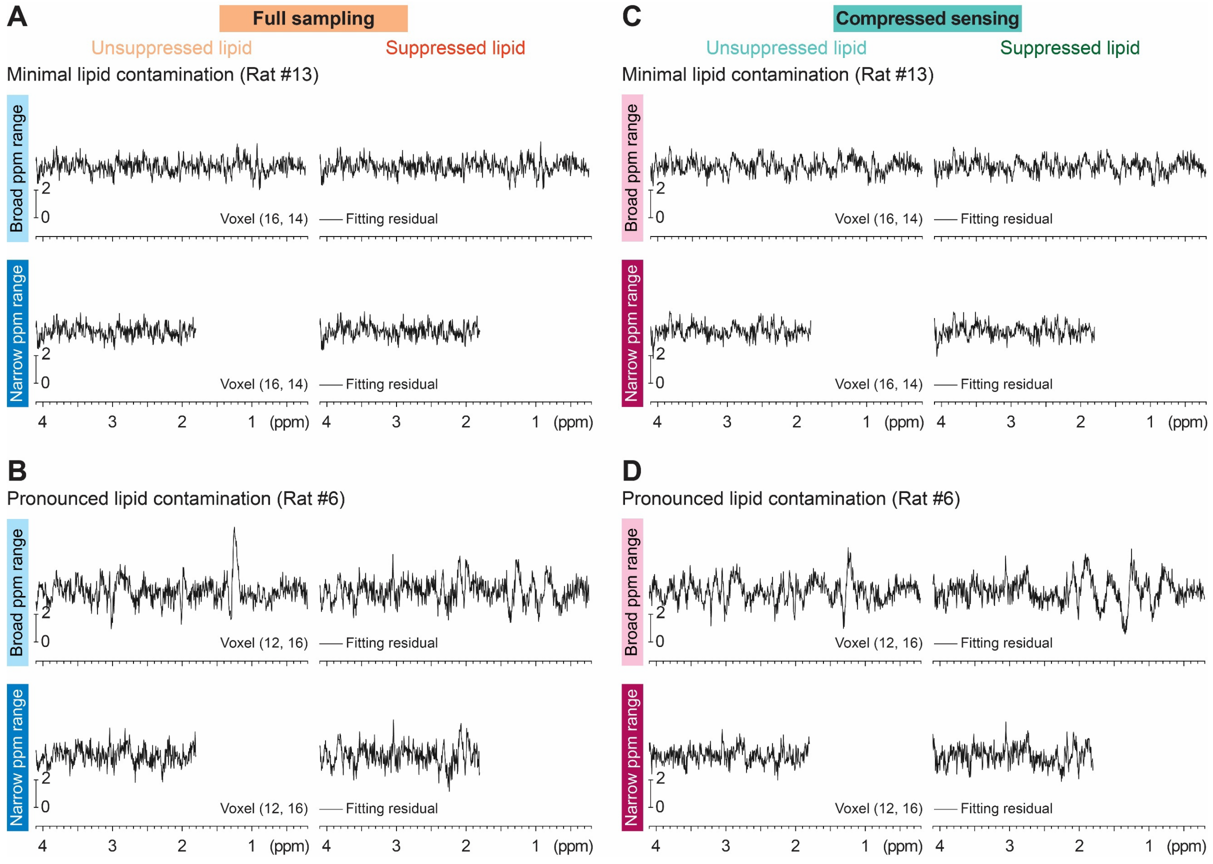


**Fig. S2. Fitting residuals under varying suppression and fitting conditions in Fig. 2**

(**A**) Fitting residuals of representative spectra from a fully sampled dataset with minimal lipid contamination (5 lipid components). Top: Fitting residuals over the broad fitting range without (left) and with (right) lipid suppression. Bottom: Fitting residuals over the narrow fitting range without (left) and with (right) lipid suppression.

(**B**) Same as (A) but with pronounced lipid contamination (23 lipid components).

(**C**) Same as (A) for a compressed sensing dataset (5 lipid components).

(**D**) Same as (B) for a compressed sensing dataset (17 lipid components).

All signal amplitudes are presented in arbitrary units.
